# STAT5-dependent regulation of CDC25A by miR-16 controls proliferation and differentiation in FLT3-ITD acute myeloid leukemia

**DOI:** 10.1101/708594

**Authors:** Gabrielle Sueur, Alison Boutet, Mathilde Gotanègre, Véronique Mansat-De Mas, Arnaud Besson, Stéphane Manenti, Sarah Bertoli

## Abstract

We recently identified the CDC25A phosphatase as a key actor in proliferation and differentiation in acute myeloid leukemia which expresses the FLT3-ITD mutation. In this paper we demonstrate that CDC25A level is controlled by a complex STAT5/miR-16 transcription and translation pathway working downstream of this receptor. First, we established by CHIP analysis that STAT5 is directly involved in FLT3-ITD-dependent *CDC25A* gene transcription. In addition, we determined that miR-16 expression is repressed by FLT3-ITD activity, and that STAT5 participates in this repression. In accordance with these results, miR-16 expression was significantly reduced in a panel of AML primary samples carrying the FLT3-ITD mutation when compared with FLT3wt cells. The expression of a miR-16 mimic reduced CDC25A protein and mRNA levels, and RNA interference-mediated down modulation of miR-16 restored CDC25A expression in response to FLT3-ITD inhibition. Finally, decreasing miR-16 expression partially restored the proliferation of cells treated with the FLT3 inhibitor AC220, while the expression of miR-16 mimic stopped this proliferation and induced monocytic differentiation of AML cells. In summary, we identified a FLT3-ITD/STAT5/miR-16/CDC25A axis essential for AML cell proliferation and differentiation.

## Introduction

Acute myeloid leukemia (AML) arises from the malignant clonal expansion of undifferentiated myeloid precursors, resulting in bone marrow failure. Although many AML patients initially respond to conventional induction therapy, relapses are common and the prognosis is very poor (1). Over the past few decades, extensive molecular characterization studies have highlighted the complex heterogeneity of AML (2) but more in-depth knowledge of the biology of AML is still required in order to rationally design more effective therapies. The most common genetic abnormalities, which occur in about 30% of AML patients, can be found within the Fms-Like Tyrosine kinase 3 (*FLT3*) gene which encodes a receptor tyrosine kinase (RTK) (2). The most frequent mutations in this gene occur via internal tandem duplication (*FLT3-ITD*) in the juxtamembrane domain, or through point mutations, usually of the Asp835 residue within the activation loop (*FLT3-TKD*). Both types of mutations result in constitutive activation of FLT3 and promote leukemic cell proliferation and survival. The poor outcome associated with FLT3-ITD mutations (3) has generated great interest in the development of specific tyrosine kinase inhibitors (TKI). Recently, an improvement in overall survival was shown for the first time through a randomized phase-3 controlled trial that combined first-generation multi-targeted type II FLT3-TKI midostaurin (PKC412) with standard chemotherapy in both newly diagnosed FLT3-ITD and FLT3-TKD AML, and which led to the approval of midostaurin in this indication in 2017 and its common use since then (4). Among the next generation of highly specific FLT3 TKIs that have been developed, quizartinib (AC220) compares favorably in monotherapy to standard treatment in refractory or relapsed AML (QuANTUM-R trial (NCT02039726) (5)), and is currently being evaluated in first line treatment in combination with chemotherapy (QuANTUM-First trial (NCT02668653)). However, resistance to FLT3 TKI treatment still commonly occurs due to the persistence of downstream signaling pathways (*e*.*g*. MAPK), point mutations (*e*.*g*. FLT3-TKD), or micro-environmental factors (3). Although significant efforts are being made to develop more efficient tyrosine kinase inhibitors (*e*.*g*. potent type II FLT3 TKI, namely gilteritinib, crenolanib or FF-10101) or to combine FLT3 inhibitors with other targeted therapies (*e*.*g*., with Bcl2 inhibitor venetoclax (6), NCT03625505), novel approaches to eradicate FLT3-mutated leukemic cells are still lacking.

Although cell signaling pathways activated by FLT3-ITD are well characterized, cell cycle proteins that may account for deregulated proliferation of these cells remain poorly documented. In a recent work, we identified the dual-specificity phosphatase CDC25A as an early target of FLT3-ITD signaling and as a master actor in proliferation and differentiation arrest of these leukemic cells (7). CDC25A is a dual-specificity phosphatase that drives cyclin-dependent kinases activation during the cell cycle, with important functions during replication, mitosis, and G1 progression. CDC25A knock-out is lethal at an early stage of embryonic development. Its overexpression has been described in different categories of cancers, and is often associated with a poor prognosis (8). However, there are only a few studies concerning CDC25A in AML or other myeloid malignancies. CDC25A expression is increased by leukemic cell adhesion to fibronectin, and it participates in the adhesion-dependent increased proliferation of these cells (9). CDC25A is also constitutively expressed downstream of oncogenic tyrosine kinases, including NPM-ALK and BCR-ABL (10), as well as JAK2V617F in myeloproliferative neoplasms (11). CDC25A is finely regulated both at the gene and protein levels (12) and moderate variations in its expression affect genomic stability and oncogenic transformation (13). Consequently, better understanding of CDC25A regulation pathways in pathophysiological models appears to be of highest interest in order to identify new potential therapeutic targets.

We recently identified the STAT5 transcription factor as an intermediate between FLT3-ITD and CDC25A regulation in AML. However, the precise mechanism of this regulation remained undefined. In this paper we demonstrate that miR-16 regulates CDC25A protein levels downstream of FLT3-ITD in a STAT5-dependent manner, and that manipulating miR-16 has a high impact on the proliferation and differentiation of AML cells.

## Results

### STAT5 regulates CDC25A mRNA and protein levels

In a previous work (7), we showed that FLT3-ITD inhibition induced rapid (2 hours) down-regulation of CDC25A mRNA and protein in the FLT3-ITD AML cell line MOLM-14, and we identified STAT5 as a regulator of CDC25A protein level in this model. Based on these data, we investigated the mechanisms by which CDC25A is regulated downstream of FLT3-ITD. Because STAT5 is a transcription factor, we first examined whether RNA interference-mediated STAT5 down-regulation could impact CDC25A mRNA level. As shown in figure 1A, 24h after transfecting MV4-11 cells with STAT5 siRNA, the level of CDC25A mRNA was decreased by 50%, and CDC25A protein level was dramatically reduced at the same time (Figure 1A). Similarly, a pharmacological inhibitor of STAT5 reduced both CDC25A mRNA and protein levels as early as 4 hours after treatment (Figure 1B).

**Figure 1:**
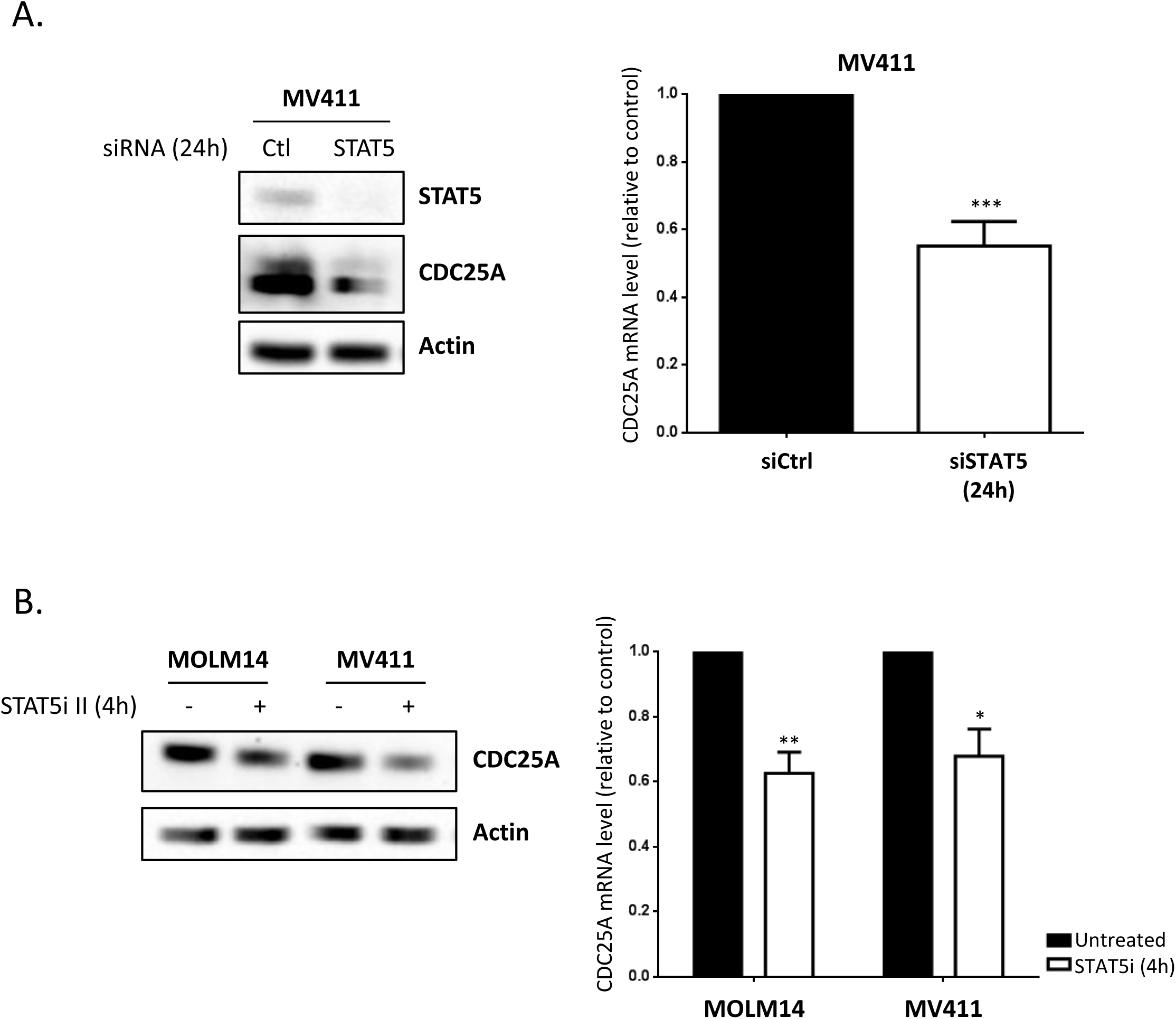
STAT5 regulates CDC25A protein and mRNA level in FLT3-ITD AML cells. A. MV411 cells were transfected for 24h (hours) with siRNA against STAT5A/B, and the CDC25A protein and mRNA levels were analyzed by western blot (left panel) and RT-qPCR (right panel) respectively. B. MOLM14 and MV411 cells were treated for 4h with STAT5 inhibitor II (1µM), and the CDC25A protein and mRNA levels were analyzed by western blot (left panel) and RT-qPCR (right panel) respectively. These results are representative of at least 3 independent experiments. Error bars represent the standard error of the mean (SEM). Actin was used as a loading control in the western blot experiments.

These data suggest that STAT5 regulates CDC25A, either directly through gene transcription, or indirectly through post-transcriptional mechanisms that affect its mRNA level, or both.

### FLT3-ITD and STAT5 are involved in *CDC25A* transcription

To establish whether FLT3-ITD activity and STAT5 modulate *CDC25A* transcription, we performed chromatin-immunoprecipitation (ChIP) experiments on MOLM-14 or MV4-11 AML cells treated with an FLT3 inhibitor. The experimental design is shown in figure 2A. First, we performed RNA Polymerase II immune-precipitation and found that the association of RNA polymerase II with the *CDC25A* promoter was disrupted in the presence of the FLT3 inhibitor (figure 2B), indicating that FLT3-ITD activity indeed controls CDC25A transcription. Similar experiments performed with a STAT5 inhibitor suggest that STAT5 is involved in this transcriptional regulation (Figure 2C). To obtain direct evidence of CDC25A transcriptional regulation by STAT5, we then performed ChIP experiments with a STAT5 antibody in MOLM-14 cells treated with the FLT3 inhibitor. These experiments revealed (i) that the STAT5 factor is indeed present on the *CDC25A* promoter in these leukemic cells, and (ii) that this association is dependent on FLT3-ITD activity (Figure 2D). To our knowledge these data represent the first direct evidence of a transcriptional regulation of CDC25A by STAT5, and they demonstrate that FLT3-ITD activity controls this process.

**Figure 2:**
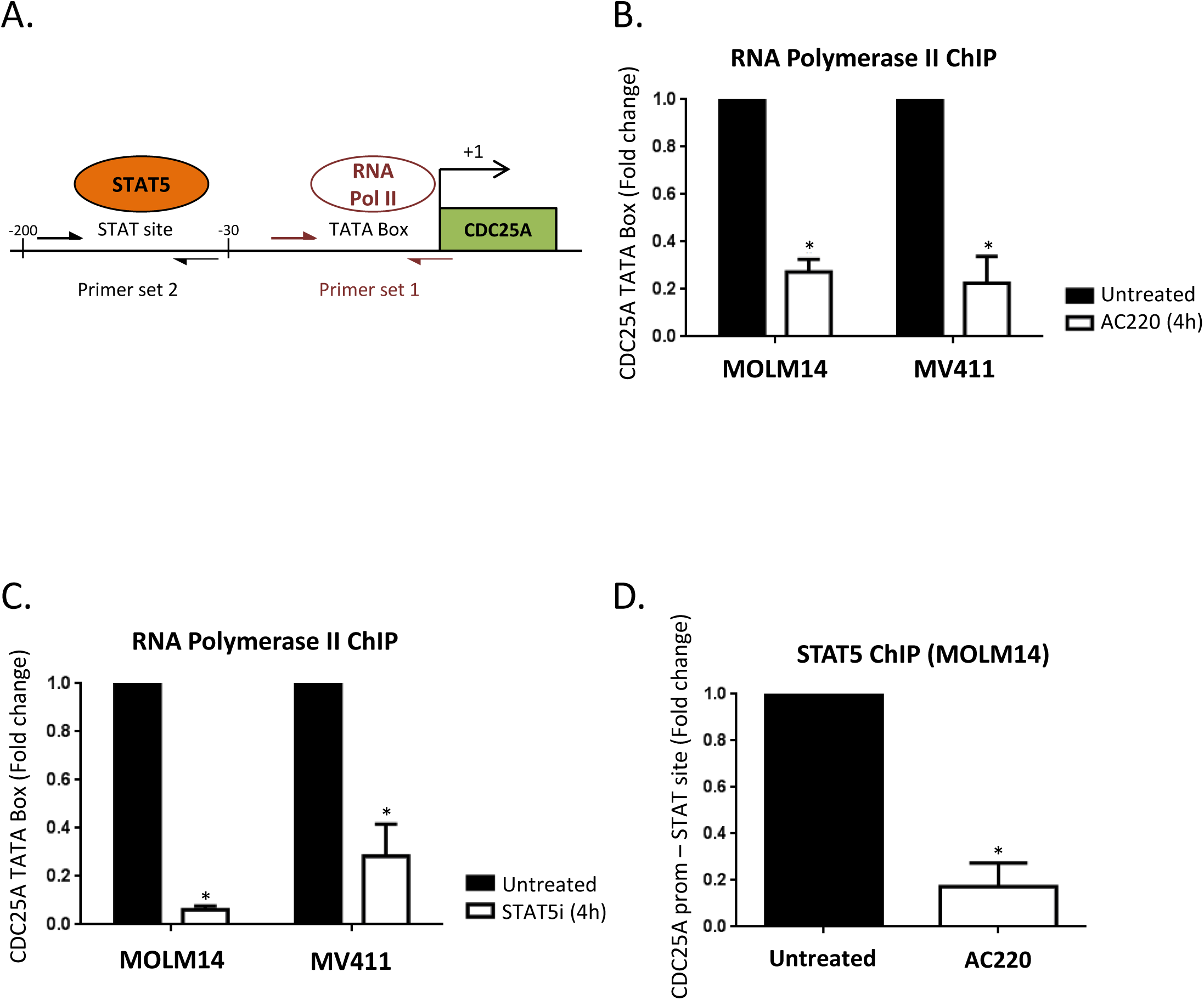
STAT5-dependent transcriptional regulation of CDC25A downstream of FLT3-ITD. A. Experimental design and position of the Q-PCR primer in the CDC25A promoter used for the CHIP experiments performed on MOLM-14 and MV4-11 cells. B. MOLM14 and MV411 cells were treated for 4h with the FLT3 inhibitor AC220 (2nM). ChIP experiments were then performed with RNA Pol II antibody and mouse IgG as a control. The CDC25A TATA Box region was amplified by qPCR as described in A. C. MOLM14 and MV411 cells were treated for 4h with STAT5 inhibitor (1 µM). ChIP experiments were then performed with RNA Pol II antibody and mouse IgG as a control. The CDC25A TATA Box region was amplified by qPCR. D. MOLM14 cells were treated for 4h with AC220 (2nM). ChIP experiments were then performed with STAT5 A/B antibody and rabbit IgG as a control. The CDC25A proximal promoter region was amplified by qPCR. These results are representative of at least 3 independent experiments. Error bars represent the SEM.

### FLT3-ITD and STAT5 induce miR-16 down-regulation in AML cells

Since CDC25A is extensively regulated at transcriptional and post-transcriptional levels by several pathways (12), we examined whether other mechanisms besides transcription could account for modifications in CDC25A level in AML cells. We specifically focused our attention on the micro RNA miR-16, a known tumor suppressor that has been reported to regulate CDC25A in response to UV-induced DNA damage (16), and which was found to be down-regulated in murine FDCP-1 cells that express FLT3-ITD (17). First, we tested the impact of FLT3 inhibition on miR-16 level in human MOLM-14 and MV4-11 cells. As shown in figure 3A, miR-16 was significantly increased after two hours of FLT3-ITD inhibition. In order to investigate the physiological meaning of these data, we established the status of miR-16 expression in primary cells from AML patients. As shown in figure 3B, miR-16 level was significantly lower in AML cells that express FLT3-ITD, further confirming that FLT3-ITD activity negatively regulates miR-16 expression. We then examined whether STAT5 could also regulate miR-16 expression in these cell lines. RNA interference-mediated down-regulation of STAT5 indeed increased miR-16 level to the same extent as FLT3 inhibition (figure 3C). In contrast, inhibiting either ERK or Akt, two other important pathways constitutively activated downstream of FLT3-ITD, did not modify miR-16 levels (figure 3D). From these experiments, we concluded that miR-16 level is negatively controlled by FLT3-ITD in a STAT5-dependent way in AML. Since miR-16 is part of two complex loci also expressing either miR-15-a, or miR-15-b, we also tested whether the level of these 2 micro-RNAs was dependent of FLT3 activity. As shown in Supplementary figure 1, FLT3 inhibition did not impact the levels of miR-15-a, or b, which indicates that the regulation of miR-16 expression by FLT3-ITD is not a regulation of either of the miR-16 cluster genes, as this would also impact miR-15-a or b expression.

**Figure 3:**
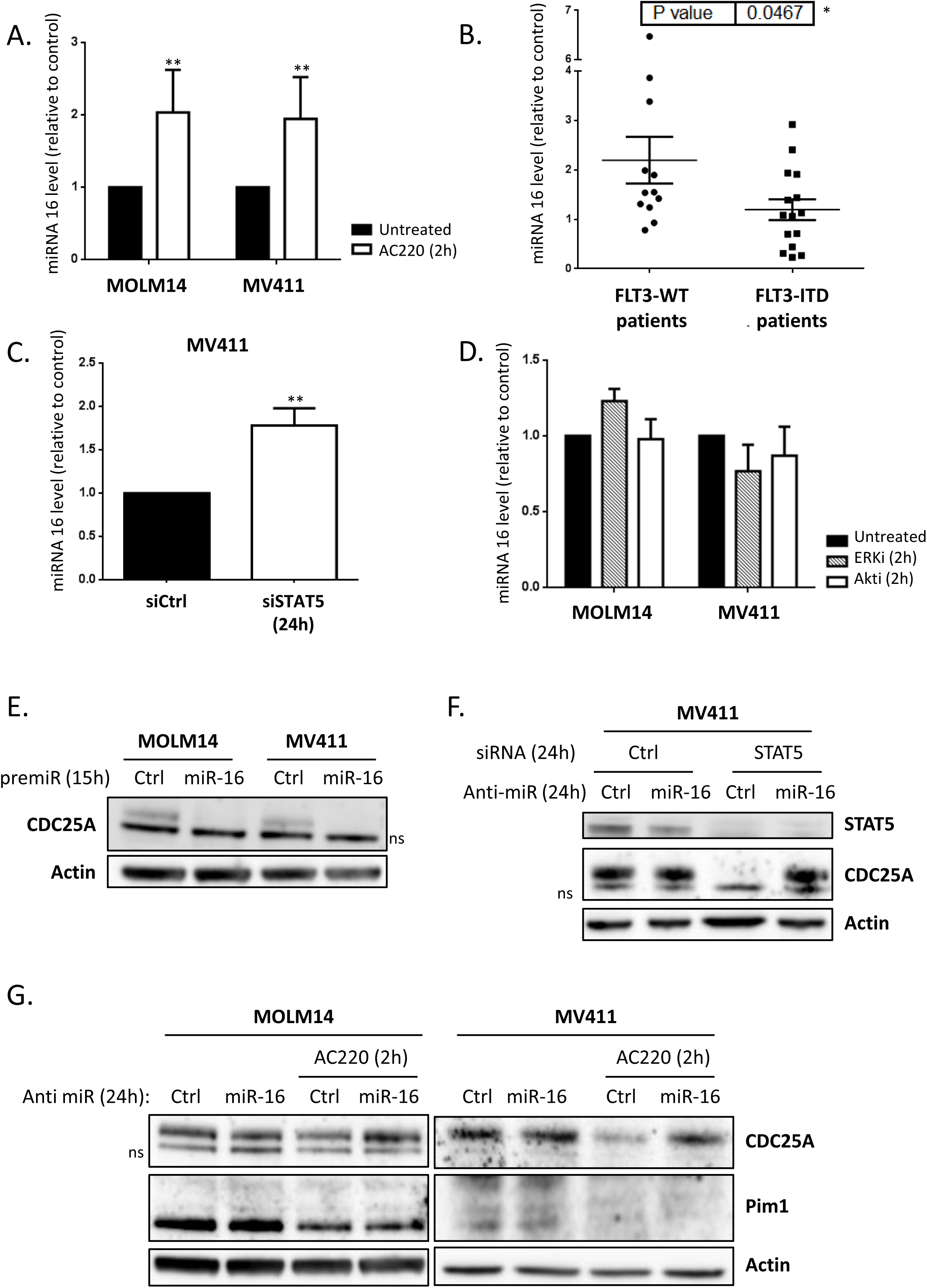
miR-16 level is dependent on FLT3-ITD activity and STAT5 in AML cells. A. MOLM-14 and MV4-11 cells were treated for 2h with the FLT3 inhibitor AC220 (2nM) and the miR-16 level was analyzed by RT-qPCR. B. Basal miR-16 levels were analyzed by RT-qPCR and normalized to a control (MOLM14). A break in the Y axis was added for easier presentation of the results. C. MV4-11 cells were transfected for 24h with STAT5A/B siRNA and the miR-16 level was analyzed by RT-qPCR. D. MOLM-14 and MV4-11 cells were treated for 2h with ERK or Akt inhibitors (50nM and 5µM respectively) and the miR-16 level was analyzed by RT-qPCR. E. MOLM-14 and MV4-11 cells were transfected for 24h with a premiR-16 and the CDC25A protein level was analyzed by western blot. F. MOLM-14 and MV4-11 cells were transfected for 24h with a miR-16 inhibitor and treated for 2hwith AC220 (2nM), the CDC25A and Pim1 protein levels were analyzed by western blot. G. MV4-11 cells were transfected for 24h with a miR-16 inhibitor and STAT5A/B siRNA, and the CDC25A protein level was analyzed by western blot. These results are representative of at least 3 independent experiments. Error bars represent the SEM. Actin was used as a loading control in the western blot experiments. ns: non-specific.

### MiR-16 regulates CDC25A protein level

We then examined whether miR-16 regulates CDC25A protein level downstream of FLT3-ITD. Transfection of FLT3-ITD AML cell lines with miR-16 mimic indeed dramatically reduced CDC25A protein levels (figure 3E). The low level of CDC25A protein in these experiments is caused by the cell cycle arrest induced after cell transfection. To further specify this process, we then used an RNA interference strategy to repress miR-16 expression in MOLM-14 and MV4-11 cells treated with the FLT3i AC220 (figure 3G). As expected, down-regulating miR-16 in these conditions totally rescued CDC25A protein expression on FLT3 inhibition, demonstrating that miR-16 is the major regulator of CDC25A protein level downstream of FLT3-ITD. Interestingly, in opposition to results obtained by Kim el al (17), we observed no rescue of the Pim1 kinase level by miR-16 inhibition, suggesting that miR-16 does not regulate Pim1 expression downstream of FLT3-ITD in our cell model. We then tested whether miR-16 also regulates CDC25A protein expression downstream of STAT5. As expected, inhibiting miR-16 completely restored CDC25A protein expression in MOLM-14 cells transfected with STAT5 siRNA (figure 3F), which confirms the existence of a FLT3-ITD/STAT5/miR-16/CDC25A axis in these AML cells.

To further specify how miR-16 regulates CDC25A translation, we then measured CDC25A mRNA level by RT-QPCR. In contrast to our observations concerning CDC25A protein level, miR-16 inhibition only very partially restored CDC25A mRNA expression when FLT3-ITD is inhibited in MOLM-14 cells, and not at all when FLT3-ITD is inhibited or STAT5 is down-regulated in MV4-11 cells (Supplementary Fig 2 A and B). Moreover, when CDC25A mRNA stability was studied using an Actinomycin-D chase experiment only a low impact of FLT3 inhibition (Supplementary Fig 2 C) and no impact of STAT5 knockdown (Supplementary Fig 2D) on CDC25A mRNA half-life were shown. Altogether these results imply that although miR-16 might somehow destabilize CDC25A mRNA, its major function resides in the inhibition of mRNA translation.

### The STAT5/miR-16/CDC25A axis is specific to FLT3-ITD AML cells

We then examined whether these molecular pathways specifically work downstream of FLT3-ITD. In order to do this, we first used the HEL leukemic cell line, which expresses the mutant tyrosine kinase JAK2V617F but not the FLT3 receptor, and in which we previously described a STAT5-dependent regulation of CDC25A (11). As shown in figure 4A, RNA interference-mediated down-regulation of STAT5 actually induced a dramatic decrease in CDC25A protein level, but inhibiting miR-16, which was modestly increased under these conditions (figure 4B), did not restore CDC25A protein level. This indicates that STAT5 regulation of CDC25A does not depend on miR-16 in this cell line. In the OCI-AML3 leukemic cell line, which expresses wild type FLT3, activating FLT3 with FLT3 ligand, or inhibiting the receptor with AC220, modified neither CDC25A protein nor the miR-16 level (Figure 4C-D). Overall, these data obtained in human leukemic cells suggest that several different signaling pathways operate in different cell lines to regulate CDC25A, and that the STAT5/miR-16/CDC25A pathway specifically acts downstream of the FLT3-ITD mutant receptor in AML.

**Figure 4:**
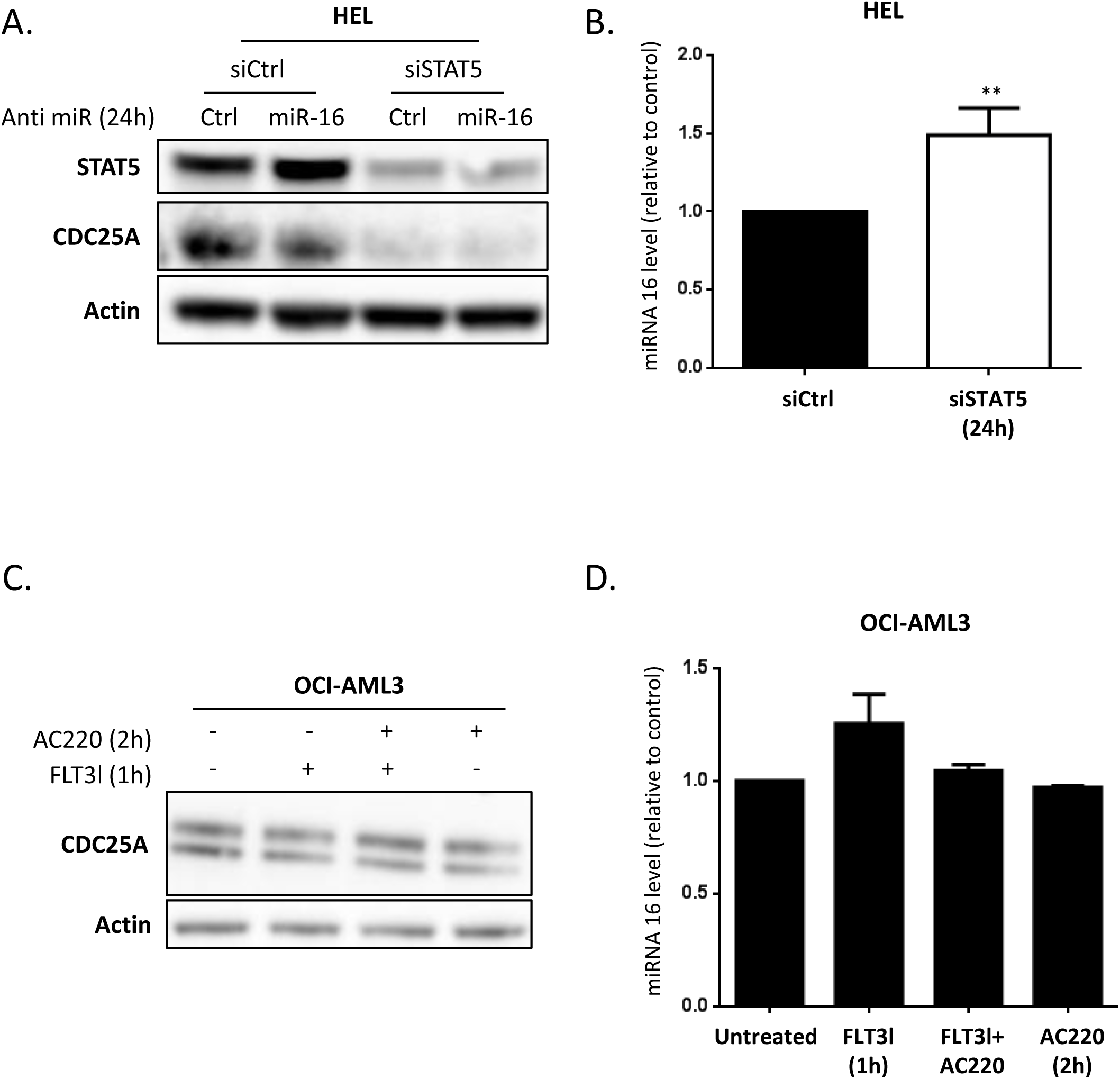
The STAT5/miR-16/CDC25A axis is specific to FLT3-ITD positive cells. A. HEL cells were transfected for 24h with a miR-16 inhibitor and STAT5A/B siRNA. CDC25A and STAT5 protein levels were analyzed by western blot. B. HEL cells were transfected for 24h with STAT5A/B siRNA and miR-16 level was analyzed by RT-qPCR. C. OCI-AML3 cells were treated for 2h with the FLT3 inhibitor AC220 (2nM) and/or FLT3 ligand (6nM), and the CDC25A protein level was analyzed by western blot. D. OCI-AML3 cells were treated for 2h with the FLT3 inhibitor AC220 (2nM) and/or FLT3 ligand (6nM) and the miR-16 level was analyzed by RT-qPCR. These results are representative of at least 3 independent experiments. Error bars represent the SEM. Actin was used as a loading control in the western blot experiments. ns: non-specific

### MiR-16 regulates FLT3-ITD AML cell proliferation and differentiation

Considering that we identified CDC25A as a master regulator of FLT3-ITD AML proliferation and differentiation, we decided to investigate whether this is also the case for miR-16. Expressing miR-16 mimic in MOLM-14 or MV4-11 cells for 72hsignificantly decreased their proliferation (figure 5A). Interestingly, this was not the case in the OCI-AML3 AML cells that express wild type FLT3. Down-regulating miR-16 partially restored proliferation of MOLM-14 or MV4-11 cells treated with the FLT3 inhibitor AC220 (Figure 5B), suggesting that miR-16 regulation and functions could be central to the therapeutic effects of FLT3 inhibitors. We then tested the importance of miR-16 in a MOLM-14-TKD cell line that is resistant to FLT3-ITD inhibition (7, 18). Overexpressing miR-16 also reduced cell proliferation in this case, suggesting that targeting this pathway could overcome some of the resistance observed with FLT3-ITD kinase inhibitors (Figure 5A). In accordance with these results, the clonogenic potential of primary cells from a FLT3-ITD-positive patient was reduced when miR-16 was overexpressed, while there was no impact in FLT3wt primary samples (Figure 5C).

**Figure 5:**
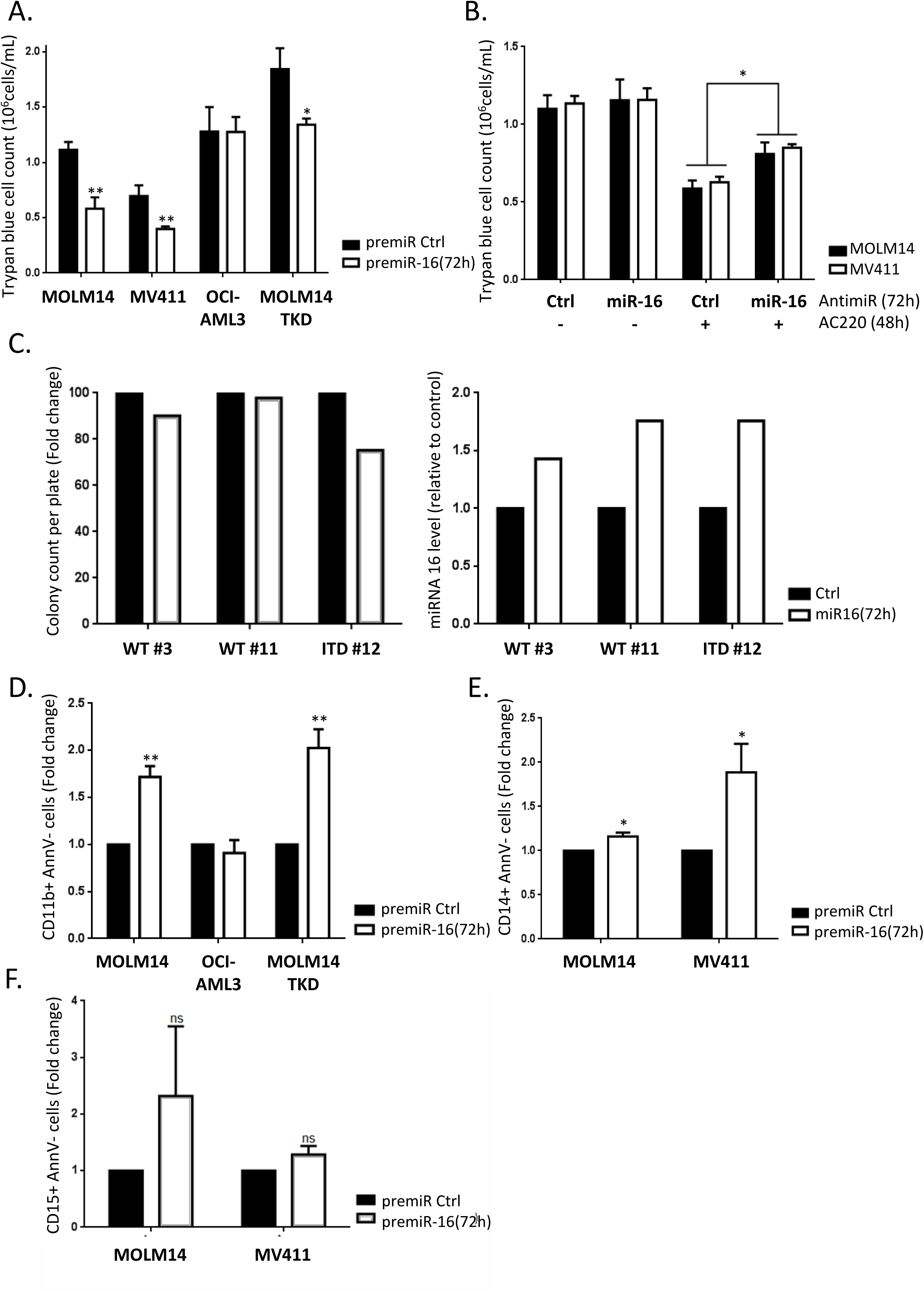
miR-16 inhibits cell proliferation in FLT3-ITD cell lines and primary cells, and triggers monocytic differentiation in FLT3-ITD AML cells. A. MOLM-14 and MV4-11 (FLT3-ITD), OCI-AML3 (FLT3wt) and MOLM-14-TKD (FLT3-ITD inhibition resistant) cell lines were transfected with pre-miRcontrol or pre-miR-16 and live cells were counted at 72h after trypan blue staining. B. MOLM-14 and MV4-11 cells were transfected with pre-miRcontrol or pre-miR-16 as in A, and then treated with AC220 (2 nM) 24h after transfection. 72h after transfection, cells were counted after trypan blue staining. C. Primary cells from three AML patients were transduced with a lentiviral vector expressing a scrambled shRNA (Ctrl, black bars) or the miR-16 (miR-16, white bars), and clonogenic potential was assessed by colony forming assay 7 days after transduction (left panel). Transduction efficiency was assessed by RT-qPCR after 3 days (right panel) D - F. MOLM-14, OCI-AML3, MOLM14-TKD or MV4-11 cells were transfected with pre-miRcontrol or pre-miR-16 as in A, and then labeled with anti CD11 (D) and CD14 (E) or anti CD15 (F) antibodies in order to monitor their differentiation state. Annexin V labeling was performed in parallel in order to exclude dying cells from the FACS analysis. These results are representative of at least 3 independent experiments. Error bars represent the SEM.

In addition to proliferation, we then examined whether miR-16 down-regulation by FLT3-ITD was involved in the differentiation block of these cells. As seen in figure 5D, miR-16 expression induced differentiation of MOLM-14 cells, as indicated by increased expression of the cell surface differentiation marker CD11b. Interestingly, this was also the case in MOLM-14-TKD cells that are resistant to FLT3 inhibition, but not in cells that express wild type FLT3 (OCI-AML3). Finally, we also observed re-expression of the CD14 but not the CD15 surface differentiation marker with miR-16 overexpression (figures 5E and F), suggesting that monocytic rather than granulocytic differentiation was induced by miR-16 in FLT3-ITD cells, which is highly consistent with our previous work on CDC25A (7).

## Discussion

In this work we investigated the molecular mechanisms that control *CDC25A* transcription and translation in acute myeloid leukemia cells that express the FLT3-ITD mutant tyrosine kinase receptor. This investigation picks up where our previous study leaves off, in which we identified this phosphatase as a master determinant of proliferation and differentiation arrest in this leukemic subtype (7). In general, CDC25A expression is tightly regulated in myeloid malignancies that express tyrosine kinase mutations (7, 11, 19) in which STAT5 and STAT3 pathways are most often activated. Transcriptional regulations of CDC25A by E2F1 (20), myc (21) and STAT3 (22) have been described previously in different cellular contexts. However, the question of STAT5-mediated transcriptional regulation of CDC25A remained an open one. In our study, STAT5 is identified at the CDC25A promoter for the first time, and we highlight the importance of STAT5 activity for proper CDC25A transcription downstream of the FLT3-ITD receptor. It has been reported that STAT3 associates with myc or Rb to respectively activate or inhibit CDC25A transcription (22). Therefore, whether STAT5 associates with transcriptional partners to finely tune CDC25A transcription in AML cells remains an interesting question.

The importance of STAT factors in hematological models has been extensively documented (23, 24), and researchers have been taking interest in the clinical relevance of these factors as targets in various cancer models (25, 26). Recently, a study in MDS and AML models showed an impairment in leukemic growth *in vivo* and an increase in differentiation in primary progenitor cells with the use of a clinical antisense nucleotide that targets STAT3 (27). Our current findings along with those of other studies (28, 29) highlight and specify the importance of STAT5 as part of an essential pathway for FLT3-ITD AML proliferation and survival, and efforts to target STAT5 in AML are in progress (30).

In addition to this direct transcriptional regulation, we have highlighted another CDC25A regulation pathway downstream of FLT3-ITD involving STAT5 as well. In fact, we identified micro-RNA 16 as a key negative regulator of CDC25A protein expression downstream of FLT3-ITD and STAT5 in AML cells. First, our data support the argument that STAT5 is an actor in miR-16 regulation downstream of FLT3-ITD. STAT5 has been shown to regulate miR-16-2 and miR15-b expression in multiple myeloma (31) by inhibiting the transcription of their cluster gene, SMC4. However, in AML cell lines, we found no effect of FLT3 inhibition on miR-15-a, or -b expression, which suggests a more complex regulation of miR-16 that may involve maturation rather than transcriptional regulation of one or both cluster genes (for a review see 32).

Our data also point to the fact that miR-16 is an important determinant of CDC25A protein level downstream of FLT3-ITD. MiR-16-dependent regulation of CDC25A has been previously reported in response to UV radiation-induced DNA damage, participating in the checkpoint response and proliferation arrest observed in these conditions (16). In this study, we demonstrate that miR-16-dependent regulation of CDC25A is not restricted to DNA damage response, and could participate in the response of leukemic cells to tyrosine kinase inhibitors. There are 2 seed sequences of miR-16 in the 3’UTR region of CDC25A mRNA. This strongly suggests a direct interaction between miR-16 and CDC25A mRNA, as was suggested with a luciferase reporter system in Hela cells in response to UV radiation (16). However, in this study we did not detect a significant impact of miR-16 expression on CDC25A mRNA level in AML cells, which suggests that depending on the cell and the environmental conditions, the functional miR-16 molecular mechanisms may differ.

In a previous work, Kim et al described a down-regulation of miR-16 by FLT3-ITD (17). Pim1 regulation by miR-16 has been described in a murine cell line model overexpressing FLT3-ITD (17). However our data do not support the argument that Pim1 is a miR-16 target in the human FLT3-ITD AML cell lines MOLM-14 and MV4-11 (see figure 3E). It remains to be established whether this discrepancy is due to the distinct AML cell models used in both studies or to other unknown parameters. However, considering that transcriptional regulation of Pim1 by STAT5 is a well-established event downstream of FLT3-ITD (33, 34), this may suggest the existence of a complex oncogenic signaling network including STAT5, miR-16, Pim1, and CDC25A that still needs to be clarified.

MiR-16 is considered to be a tumor suppressor because it negatively regulates the expression of pro-oncogenic proteins involved in cell proliferation and cell death. Therefore, some studies aimed at expressing miR-16 as a therapeutic approach have been developed in a murine model of CLL (35, 36). However the use of micro RNAs as therapeutic tools remains rather unsuccessful as they have never progressed beyond a phase I trial (37). This emphasizes the need for a clearer understanding of the networks of proteins regulated downstream of these oligonucleotides. Although we describe CDC25A as an important target of miR-16 in this study, it is unlikely that all the effects of miR-16 expression in FLT3-ITD cells are only due to CDC25A regulation. For instance, in terms of the effects of miR-16 expression on the differentiation process, it should be noted that this RNA is involved in retinoic acid induced differentiation (38), and that PU.1, an important transcriptional regulator of myeloid differentiation, has been described as a miR-16 target (39). Whether or not the miR-16-dependent differentiation that we observed in FLT3-ITD AML cells is linked with these pathways remains to be investigated. With regards to its cancer relevance, another interesting miR-16 target that has been established is the anti-apoptotic protein Bcl2 (40). Bcl2 inhibitors are currently being introduced as potential therapeutic tools in AML (41), and interestingly, a synergy between venetoclax and the FLT3-inihibitor quizartinib has been described (42). It would be interesting to establish whether miR-16-dependent regulation of Bcl2 is involved in this process.

## Methods

### Cell lines and reagents

Human acute myeloid leukemia cell lines MOLM-14, MV4-11, OCI-AML3, HEL (DSMZ, Braunschweig, Germany), and MOLM-14-TKD (provided by Dr. J Tamburini, Paris) were cultured in RPMI 1640 medium (Gibco, Life Technologies, Carlsbad, CA, USA), supplemented with 10% Fetal Bovine Serum (Gibco, Life Technologies, Carlsbad, CA, USA). All cells were cultured at 37°C and 5% CO_2_ in a humidified incubator, and were regularly tested for mycoplasma contamination. The Akt inhibitor VIII and STAT5 inhibitors were purchased from Calbiochem (San Diego, CA, USA). The FLT3 inhibitor quizartinib AC220 and the MEK inhibitor PD0325901 were purchased from Selleck Chemicals (Houston, TX, USA). FLT3 ligand was purchased from R&D Systems (Minneapolis, MN, USA), and D Actinomycin was purchased from Sigma (Saint-Louis, MO, USA).

### Western Blot

1 × 10^6^ cells were lysed in 60 μl of NuPAGE^®^ LDS Sample Buffer (Novex, Life Technologies, Carslbad, CA, USA), sonicated for 4×7 seconds, and boiled for 3 minutes. Proteins were then resolved on NuPAGE^®^ 4-12% Bis-Tris Gels and transferred to nitrocellulose membrane. The membrane was saturated for 1 hour in Tris Buffer Saline with Tween 0.1% (TBS-T) containing 5% non-fat milk. Membranes were blotted with suitable antibodies overnight at 4°C, washed thrice with TBS-T and incubated for 1 hour with HRP-conjugated secondary antibody (Promega, Madison, WI, USA). After four additional washes, detection was achieved with ECL™ or ECL™ prime western blotting detection reagents (GE Healthcare, Chicago, IL, USA). The antibodies used were: monoclonal anti-CDC25A (F-6, sc-7389) and anti-Pim1 (12H8, sc-13513) from Santa Cruz; polyclonal anti-phospho-STAT5 A/B (Tyr 694, 9351), anti-STAT5 A/B (9363), from Cell Signaling Technology (Beverly, MA, USA) and anti-α-actin (MAB1501) from Millipore (Burlington, MA, USA).

### RT-qPCR

Cells were lysed in TRIzol (Invitrogen, Life Technologies, Carlsbad, CA, USA) and total RNA was then extracted according to the manufacturer’s instructions. For gene expression analysis, cDNA was generated with the SuperScript III First-Strand Synthesis System for RT-PCR (Invitrogen, Life Technologies, Carlsbad, CA, USA) according to the manufacturer’s instructions. The PCR was performed with TaqMan^®^ Gene Expression Master Mix (Applied Biosystems, Life Technologies, Carlsbad, CA, USA) with 10ng of cDNA, The primer used was Hs00947994_m1 (Applied Biosystem, Life Technologies, Carlsbad, CA, USA) for *CDC25A. GUSB* (Hs00939627_m1) and *B2M* (Hs00984230_m1) were used as housekeeping genes. For mature micro-RNA expression analysis, cDNA was generated using the TaqMan^®^ micro-RNA reverse transcription kit (Applied Biosystems, Life Technologies, Carlsbad, CA, USA), and PCR was performed using the TaqMan^®^ Universal Master Mix with UNG (Applied Biosystems, Life Technologies, Carlsbad, CA, USA), and the TaqMan microRNA assays (Applied Biosystems, Life Technologies, Carlsbad, CA, USA) according to the manufacturer’s instructions. The primers used were miR-16 (assay 000391) and U6 snRNA (assay 001973) as a control. Results were analyzed with the StepOne™ software v2.2.2 (Applied Biosystems, Life Technologies, Carlsbad, CA, USA) using the conventional ΔΔCt method. PCR was carried out on a StepOne™ (Applied Biosystems, Life Technologies, Carlsbad, CA, USA).

### RNA transfection

Leukemic cell lines were transfected with the Neon transfection system (Life Technologies, Carlsbad, CA, USA). Cells (2 × 10^6^) were re-suspended in 100 μl of resuspension buffer, and various doses of RNA were added. RNA sequences were: specific STAT5A and STAT5B siRNA (ON-TARGETplus SMARTpool, human STAT5A and STAT5B, Dharmacon, Lafayette, CO, USA) or siRNA control (sigenome control pool non targeting #2, or ON-TARGETplus control pool, Dharmacon, Lafayette, CO, USA), anti-miR control or anti-miR-16 (Life Technologies, Carlsbad, CA, USA), and pre-microRNA negative control or pre-microRNA-16 (Life Technologies, Carlsbad, CA, USA). Cells were then transfected with the nucleofector device (program 5 for MOLM14, MV4-11 and OCI-AML3 cell lines, custom program for the HEL cell line), and subsequently re-suspended in normal culture medium at a concentration of 1 × 10^6^ cells/ml.

### Chromatin immuno-precipitation (ChIP)

Cells were crosslinked, lysed and sonicated according to (14), then STAT5 and RNA polymerase II immunoprecipitation was performed according to (15). The antibodies used for immunoprecipitation were STAT5 A (sc-1081, Santa Cruz Biotechnology, Santa Cruz, CA, USA) and STAT5 B (sc-835, Santa Cruz Biotechnology), 1.2 µg each, and RNA Polymerase II CTD (MABI0601, Clinisciences, Nanterre, France), 2µg. DNA from ChIP was isolated with a phenol/chloroform extraction, and used for qPCR with the primers listed in Supplementary Table 1. Results were then normalized using the percent input method.

### Patient samples

Patient AML samples used in this work were obtained after informed consent in accordance with the Declaration of Helsinki and were stored in the HIMIP collection. In compliance with French law, the HIMIP collection was declared to the Ministry of Higher Education and Research (DC 2008-307 collection 1) and a transfer agreement was obtained (AC 2008-129) after approval by the ethics committees (Comité de Protection des Personnes Sud-Ouest et Outremer II and the APHP ethics committee). Clinical biological annotations of the samples were declared to the CNIL (French Data Protection Authority). All were diagnosed at the Department of Hematology of Toulouse University Hospital. The patients’ characteristics are listed in Supplementary Table 2.

### Patient samples and clonogenic assays

Patient samples’ origins and characteristics are described in the *Online Supplementary Methods* and Supplementary Table 2.

Frozen patient cells were thawed in IMDM medium with 20% FBS and immediately processed for treatment. For clonogenic assays, cells were grown in duplicate in H4230 methylcellulose medium (Stem Cell Technologies, Vancouver, BC, Canada) supplemented with 10% 5637-conditionned medium in 35mm petri dishes for 7 days in a humidified CO2 incubator. Leukemic colonies were then scored under an inverted microscope.

### Transduction of primary cells

To generate lentiviral vectors that express the micro-RNA 16, the miR-16 hairpin sequence was cloned into the pLKO.1 lentiviral vector. We used 293T packaging cells co-transfected with lentiviral protein (GAG, POL, and REV) that encode plasmids and our control (a scrambled shRNA hairpin sequence) or miR-16 plasmids. Supernatants containing lentivirus were collected for the next two day. Patient cells were thawed, plated at 1.5 million cells/ml and spinoculated for an hour with the concentrated lentiviral particles, before being inoculated for clonogenic assays.

### Flow cytometry

Cells were stained for 30 min (minutes) with the anti-human monoclonal antibody CD11b-PE (IM2581U, Beckman Coulter, Brea, CA, USA), CD14-FITC (555397, BD Biosciences, San Jose, CA, USA), or CD15-APC (551376, BD Biosciences, San Jose, CA, USA). Annexin V-FITC or APC (BD Biosciences, San Jose, CA, USA) and 7-AAD (Cell viability assay, BD Biosciences, San Jose, CA, USA) were used to assess cell viability. Data were collected on a MacsQuant cytometer (BD Biosciences, San Jose, CA, USA), and analyzed using the FlowJo software (BD Biosciences, San Jose, CA, USA)

### Statistics

Experiments in cell lines were performed at least 3 times. Results are expressed as mean value +/- SEM. Statistical analysis of the data was performed by the Mann-Whitney U test using the GraphPad Prism software, version 5.0 (GraphPad Software Inc., La Jolla, CA, USA). Differences were considered as significant for p values < 0.05 (*), p < 0.01 (**), p<0.001 (***).

## Data availability

All data generated or analysed during this study are included in this published article (and its Supplementary Information files).

### Acknowledgements

We would like to sincerely thank Laure Guitton-Sert for her technical input in designing ChIP experiments. We thank Carine Valle at the CRCT molecular biology facility and Manon Farcé at the CRCT cytometry facility for their helpful discussions of the analyses. We would also like to thank Véronique De Mas, Cécile Demur and Eric Delabesse for the management of the BRC-HIMIP Biobank (Biological Resources Centres-INSERM Midi-Pyrénées “Cytothèque des hémopathies malignes”). Gabrielle Sueur is a recipient of the Ligue Nationale contre le Cancer.

## Author contributions

G.S., S.M., and S.B. concepted and designed the study; G.S., A.Bo., and M.G. performed experiments; S.B. and V.MDM. contributed clinical samples; G.S., S.M., and S.B. wrote the manuscript; A.Be., and S.M. insured administrative, technical and material support; S.M. and S.B. supervised the project.

## Competing Interest

The authors declare no competing financial interests

## Supplementary material

Supplementary information is available at Scientific Reports’ website.

