## Supplementary figures and images for "STAT5-dependent regulation of CDC25A by miR-16 controls proliferation and differentiation in FLT3-ITD acute myeloid leukemia"

### Supplementary figure 1

**Supplementary figure 1**

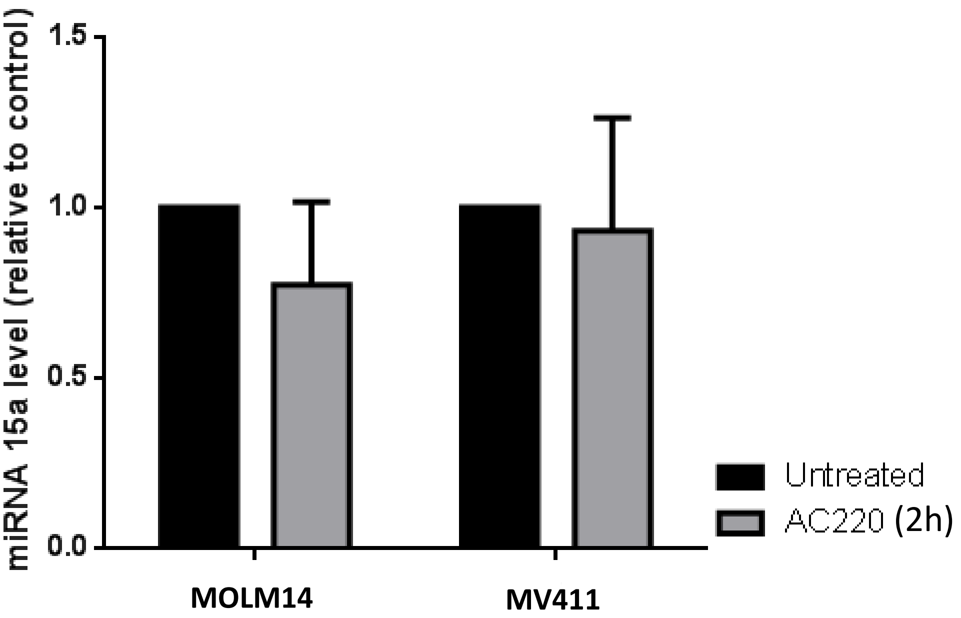


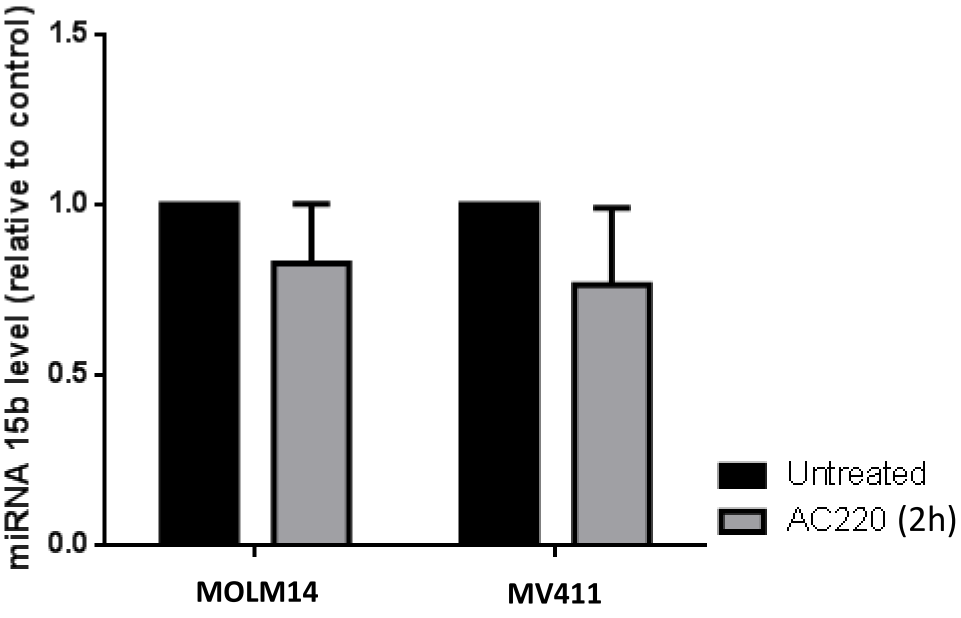

### Supplementary figure 2

#### Supplementary figure 2

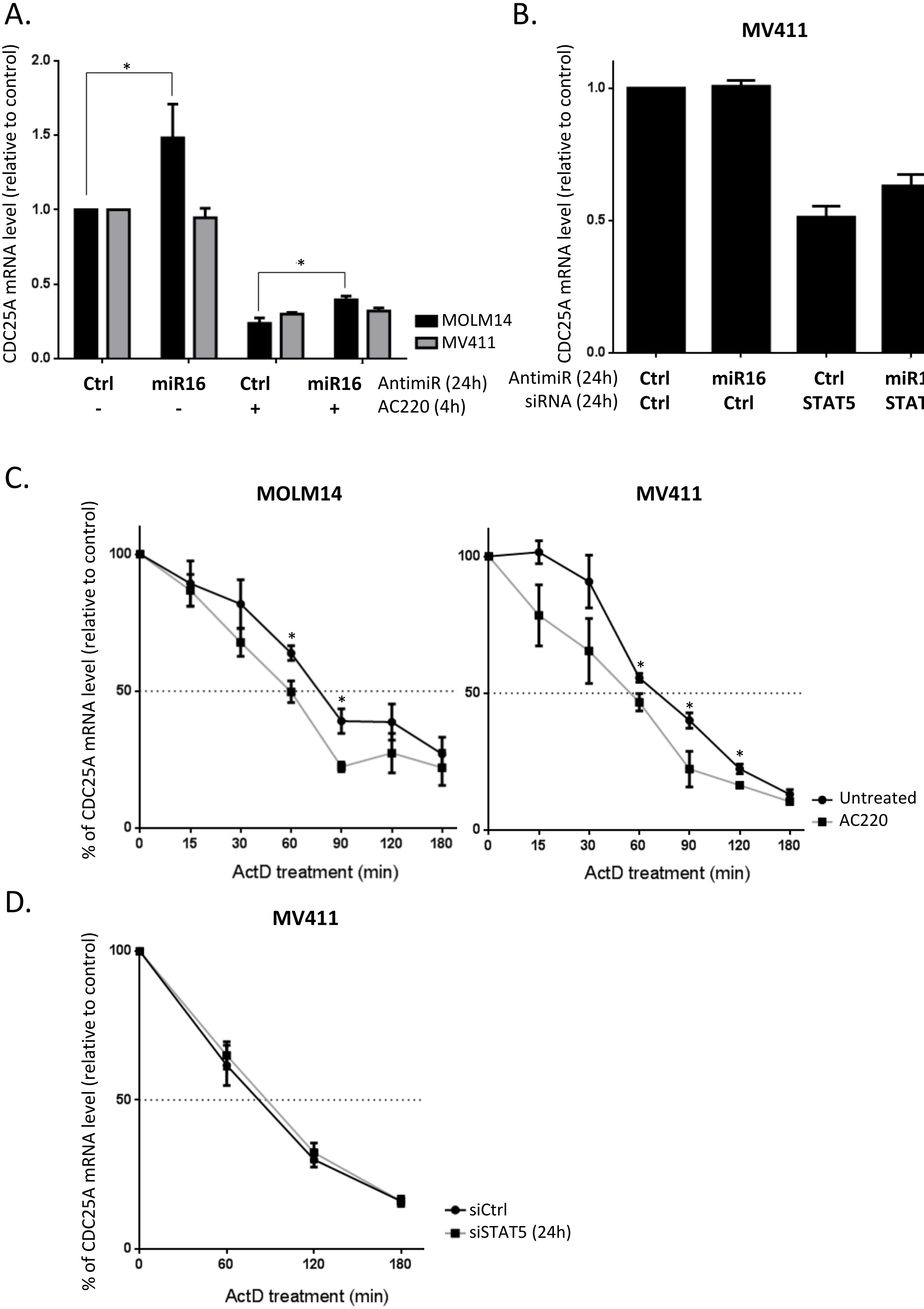
