## Supplementary figures legends for "STAT5-dependent regulation of CDC25A by miR-16 controls proliferation and differentiation in FLT3-ITD acute myeloid leukemia"

**Supplementary figure 1:** FLT3 inhibition does not modulate miR-15-a or b level

MOLM14 and MV411 cells were treated for 2h with AC220 (2nM) and miR-15-a (upper panel) and miR-15-b (lower panel) levels were analyzed by RT-qPCR. These results are representative of at least 3 independent experiments. Error bars represent the SEM.

**Supplementary figure 2**: miR-16 tenuously regulates CDC25A mRNA level

A. MOLM14 (black bars) and MV411 (grey bars) cells were transfected for 24 hours with a miR-16 inhibitor and treated for 2h with AC220 (2nM). The CDC25A mRNA level was analyzed by RT-qPCR

B. MV411 cells were transfected for 24 hours with a miR-16 inhibitor and a STAT5A/B siRNA. The CDC25A mRNA level was analyzed by RT-qPCR

C. MOLM14 and MV411 cells were treated for 1h with AC220 (2 nM) and further treated with D actinomycin (3µg/ml) for the indicated times. The CDC25A mRNA level was analyzed by RT-qPCR.

D. MV411 cells were transfected for 24h with STAT5A/B siRNA and treated with D actinomycin (3µg/ml) for the indicated times. The CDC25A mRNA level was analyzed by RT-qPCR.

These results are representative of at least 3 independent experiments. Error bars represent the SEM.
